# MAP4Ks inhibition promotes retinal neuron regeneration from Müller glia in adult mice

**DOI:** 10.1101/2022.10.14.512328

**Authors:** Houjian Zhang, Yuli Guo, Yaqiong Yang, Jingbin Zhuang, Yuqian Wang, Jiankai Zhao, Rongrong Zhang, Jiaoyue Hu, Wei Li, Jianfeng Wu, Haiwei Xu, Steven J. Fliesler, Yi Liao, Zuguo Liu

## Abstract

Mammalian Müller glia (MG) possess limited regenerative capacities. However, the intrinsic capacity of mammalian MG to transdifferentiate to generate mature neurons without transgenic manipulations remains speculative. Here we show that MAP4K4, MAP4K6 and MAP4K7, which are conserved Misshapen subfamily of ste20 kinases homologs, repress YAP activity in mammalian MG and therefore restrict their ability to be reprogrammed. However, by treating with a small molecule inhibitor of MAP4K4/6/7, mouse MG regain their ability to proliferate and enter into a retinal progenitor cell (RPC)-like state after NMDA-induced retinal damage; such plasticity was lost in YAP knockout MG. Moreover, spontaneous trans-differentiation of MG into retinal neurons expressing both amacrine and retinal ganglion cell (RGC) markers occurs after inhibitor withdrawal. Taken together, these findings suggest that MAP4Ks block the reprogramming capacity of MG in a YAP-dependent manner in adult mammals, which provides a novel avenue for the pharmaceutical induction of retinal regeneration *in vivo*.

## INTRODUCTION

Loss of photoreceptors and retinal ganglion cells (RGCs) results in vision loss in prominent retinal diseases, such as age-related macular degeneration, diabetic retinopathy, and glaucoma. Neither photoreceptors nor RGCs possess endogenous regenerative capabilities, hence, presenting a major impediment to developing cures for these blinding diseases. Regenerative medicine approaches have been tried to solve this problem, *e.g*., by transplanting stem/progenitor cells or their derived cells into the retina, but the outcomes have been variable and, in general, have not been effective ^1^. Besides, some cold-blooded vertebrate species, such as fish and amphibians, are able to regenerate retinal neurons from endogenous MG ^2–4^, which has inspired new ideas about how to repair damage to retinal neurons caused by various genetic, environmental, or traumatic insults.

Although unable to regenerate retinal neurons, mammalian MG retain some stem/progenitor cell characteristics *in vitro* ^5^ and *in vivo* ^6^. Even more attractive, early studies demonstrated that mouse MG express the cell cycle gene CyclinD3 ^7^ and upregulate the retinal progenitor cell (RPC) marker Pax6 ^8^ in response to retinal injury, which closely mimics the activation pattern observed in fish MG. Later, one landmark study proved the regenerative potential of mammalian MG by intravitreal injection of growth factors, which converts MG into amacrine cells ^9^. Recently, both gain-of-function and loss-of-function methods have successfully converted mouse MG into retinal neurons, including photoreceptors ^10^, bipolar/amacrine cells ^11–13^, as well as RGCs ^14–16^ in various mouse models. Despite these achievements, however, all these studies involved manipulation of gene expression using transgenic strategies, which has largely limited their clinical translational potential. In addition, since exogenous transcription factors were introduced, it remains an open question whether mammalian MG retain the ability to transdifferentiate into photoreceptors and RGCs.

Without growth factors or exogenous genes, mammalian MG only temporarily express cell cycle genes and the Pax6 gene in response to retinal damage, so reactive gliosis ensues within a day or so. Therefore, it is reasonable to suspect that endogenous blocking mechanisms exist that can repress regenerative capabilities of MG. Recently, cross-species sequencing has revealed that the Hippo pathway plays a pivotal role in restoring reactive MG to a quiescent state after retinal injury in mice ^13^. Specifically, conditional deletion of Yes associated protein 1(YAP1, also known as YAP, a transcriptional regulator of cellular proliferation and apoptosis) ^17,18^ in MG, a key downstream effector protein of the Hippo pathway, was shown to prevent the upregulation of cell-cycle entry genes in reactive MG in both NMDA- and MNU-damaged retina ^19,20^. Moreover, over-activation of YAP has been shown to be able to drive MG into a progenitor-like state after retinal injury in mice ^20^. All these lines of evidence indicate that the Hippo pathway could be a druggable target for retinal neuron regeneration in mammals.

The Hippo signaling pathway was initially identified in model organism *Drosophila*, which plays fundamental roles in organ size regulation, development, tissue regeneration and tumorigenesis ^17^. In the canonical Hippo pathway, Ste-20 family kinase Mst1/2 phosphorylates and activates NDR family kinase Lats1/2 and cofactor MOB1/2, which subsequently phosphorylates and sequesters YAP/TAZ in the cytoplasm. However, one intriguing finding is that Lats1/2 can still be phosphorylated in both Mst1/2-null mouse embryonic fibroblasts (MEFs) and hepatocellular carcinoma cells (HCCs) ^21^. As a result, multiple studies have demonstrated that mitogen-activated protein kinase kinase kinase kinase (MAP4K) family members, particularly HPK/GCK-like kinase (HGK/MAP4K4), misshapen-like kinase 1 (MINK1/MAP4K6), and Nck-interacting kinase (TNIK/MAP4K7) can work in parallel with Mst1/2 to phosphorylate Lats1/2 and regulate YAP/TAZ activities ^22,23^. However, until now, most studies regarding MAP4Ks-mediated YAP regulation have been conducted using mammalian cell lines, and its roles in normal and pathological cellular physiology are largely unknown or yet to be verified.

Thus, we hypothesized that MAP4Ks might block mammalian MG regenerative capabilities by suppressing the YAP/TAZ activities. Accordingly, by using a highly selective small-molecule compound, DMX-5804, which simultaneously inhibits MAP4K4/6/7 and inhibits cell death and augments mitochondrial function and calcium cycling ^24^, we showed that MAP4K4/6/7 converge to inhibit YAP activity in human and mouse MG. Notably, inhibition of MAP4K4/6/7 kinase activities in the early stage of retinal injury promoted MG proliferation and transformation into an RPC-like state by activating YAP. Even more exciting, long-term tracing experiments demonstrated the intrinsic ability of mammalian MG to regenerate amacrine and RGC-like neurons after NMDA-induced inner retinal neuron injury, which opens a new window for future advances in retinal regenerative medicine.

## RESULTS

### MAP4K4/6/7 are Expressed in Mammalian MG

First, the mRNA and protein expression levels of MAP4K4/6/7 isoforms were measured in adult murine retinas (~ 8 wk old) injured by intravitreal injection of N-methyl-D-aspartate (NMDA); age-matched mice were injected with same volume of vehicle (PBS) alone to serve as controls. Compared with PBS controls, we observed a progressive increase of all three kinases in both mRNA (Figure 1A) and protein (Figure 1B,C) levels beginning from 12 h after NMDA injection, and their expression levels were maximal by 24 h after NMDA injection. Elevated expression levels of these three kinases persisted in the neuroretina for the next 3 days (Figure 1B,C). In addition, the phosphorylation of YAP in the neuroretina after NMDA treatment was evaluated by Western blot analysis, and a similar pattern of elevated expression level was observed for both MAP4K4/6/7 and YAP phosphorylation (Figure 1B,C), which is a harbinger of a possible relationship between MAP4Ks and YAP activity.

Next, immunostaining was performed to determine the localization of MAP4K4/6/7 isoforms in the adult murine retina. As revealed by Immunofluorescence, MAP4K4/6/7 isoforms were present in all of the histological layers and all of their cell types in the neural retina (Figure 1D-G). Co-localization analysis showed that MAP4K4/6/7 were expressed in the same cells as YAP (see Figure 1H-K), a critical transcription cofactor downstream of the Hippo pathway. Previously, YAP was reported to be specifically expressed in murine MG ^25^. Therefore, these results confirmed that MAP4K4/6/7 were co-expressed with YAP in mature mouse MG, and indicated a possible regulatory role of MAP4Ks for YAP activity.

### MAP4K4/6/7 Work Redundantly to Regulate YAP Phosphorylation in Mammalian MG

Considering the co-localization and similar expression trends of MAP4K4/6/7 and YAP phosphorylation in NMDA-injured murine retina, we next tried to determine whether MAP4Ks regulated YAP activity in MG. MAP4K4/6/7 were expressed in the human MG cell line MIO-M1, which expresses MG markers glutamine synthetase (GS) and glial fibrillary acidic protein (GFAP) (Figure S1A). Among MAP4Ks, MAP4K4 was shown to have the strongest ability to regulate YAP activity, while MAP4K6/7 work in concert with MAP4K4 in HEK293A cells ^22^. Although knockdown of MAP4K4 exhibited a trend toward being able to affect YAP phosphorylation in MG, but the effect was not statistically significant (Figure S1 B-D). In contrast, when MAP4K4/6/7 were targeted by siRNAs simultaneously, we observed a ~50%, statistically significant decrease of YAP phosphorylation (Figure S1 E, F, *p*=0.0004) and an increase in YAP nuclear localization (Figure S1 G, H [*p*=0.0015], I [*p*=0.0019]). Taken together, these results suggested MAP4K4, MAP4K6 and MAP4K7 work redundantly to regulate YAP phosphorylation in MG.

### DMX-5804 Suppresses MAP4K4/6/7-mediated YAP Phosphorylation in Mammalian MG

Next, we hypothesized that pharmacological suppression of MAP4K4/6/7 could alleviate YAP phosphorylation and promote the nuclear localization of YAP in MG. Accordingly, a small-molecule inhibitor DMX-5804, which exhibits high potency and selectivity towards MAP4K4/6/7 ^24^, was utilized to suppress YAP phosphorylation in MG *in vitro* and *in vivo*. Indeed, the application of DMX-5804 in culture media significantly reduced YAP phosphorylation in MIO-M1 cells in a dose- and time-dependent manner (Figure S2A-D, *p*<0.05). Also, YAP nuclear translocation was observed when MIO-M1 cells were treated with DMX-5804 (Figure S2E-G). Because it was suggested that MAP4K4 exerts the largest contribution among MAP4Ks to regulate YAP activities, we overexpressed MAP4K4 in MIO-M1 cells and examined YAP phosphorylation after DMX-5804 treatment, so as to verify the drug specificity. As expected, the phosphorylation level of YAP was reduced at 6 h after cells were treated with DMX-5804, and the over-expression of MAP4K4 significantly reversed the inhibitory effects of DMX-5804 at various concentrations (Figure S2H-K, *p*=0.0018).

Next, we assessed the effect of DMX-5804 *in vivo*. A dose-escalating assay showed that DMX-5804 blocked YAP phosphorylation in the neuroretina at 6 h after intraperitoneal (*i.p*.) injection (dose range: 0.5 mg/kg to ~ 10 mg/kg) in comparison with the solvent control (Figure S3A-D). In addition, a single *i.p*. injection of 2 mg/kg DMX-5804 could suppress YAP phosphorylation by ~ 40% for more than 6 h in the intact adult murine retina (Figure S3E, F, *p*<0.05). Immunofluorescence staining further verified that DMX-5804 treatment promoted YAP nuclear translocation in MG, which was assessed by the co-localization with MG nuclear markers Sox9 (Sox9+ MG) (Figure S3G-I). Importantly, these results suggested that the lipid-soluble small molecule DMX-5804 possesses the ability to cross the blood-retinal barrier to regulate YAP activities in MG when drug is administrated systemically (*i.e*., by *i.p*. injection).

To assess the effect of DMX-5804 in the NMDA-injured retina, we developed a protocol for continuous *i.p*. injection (4 injections per day, from 6 h post-NMDA injection until 5 days post-NMDA injection) in a mouse model with NMDA-induced retinal degeneration (Figure 2A). A progressive increase in YAP phosphorylation was observed in the NMDA-damaged neuroretina, which was significantly reduced after systemic DMX-5804 treatment (Figure 2B-E, *p*=0.0001). Moreover, more YAP was translocated into the nucleus of MG in DMX-5804 treated mice (Figure 2F-I). Taken together, these data indicate that targeting MAP4K4/6/7 activated YAP and promoted YAP nuclear translocation in reactive MG.

### DMX-5804 Promotes MG Proliferation in the NMDA-Injured Retina

In previous studies, mice carrying a constitutively active YAP5SA transgene possess more EdU^+^ MG and more cells reside in a progenitor-like state after retinal damage ^20^. Thus, we assumed that suppression of MAP4K4/6/7 pharmacologically may promote MG proliferation and dedifferentiation into a progenitor-like state. To test this hypothesis, we assessed MG proliferation using EdU labeling when DMX-5804 was injected into NMDA-treated mice following a protocol we depicted in Figure 3A. Consistent with previous studies ^8,26^, we failed to detect EdU^+^ MG, which were marked by nuclear staining of Sox9, in control (non-treated, NT), NMDA-treated or NMDA/solvent control groups (Figure 3B). In contrast, DMX-5804 treatment promoted a number of reactive MG to proliferate on the 3^rd^ day after NMDA-induced damage, as shown by EdU^+^ labeling in SOX9^+^ cells (Figure 3B). Moreover, we observed double-positive (EdU^+^/GFAP^+^) cells after DMX-5804 treatment in the injured retina, suggesting that MG in the adult murine retina might undergo a reprogramming event accompanying with a gliotic response (characterized by MG hypertrophy and increased GFAP expression) similar to zebrafish (Figure 3C). Close examination revealed that these proliferating cells exhibited similar morphologies as clonally expanding zebrafish progenitor-like cells derived from MG ^27,28^ (Figure 3C).

To evaluate DMX-5804-promoted MG proliferation in detail, mice were treated following the protocol described in Figure 2A. Subsequently, EdU was *i.p*. injected 24 h before the collection of retinal samples (Figure 3D). It is worthy to notice that a few EdU^+^/SOX9^+^ MG began to appear at 2 dpi, and this population reached the maximum number at 3 dpi. However, the number of double-positive cells was dramatically decreased at 4 dpi and 5 dpi despite continuous DMX-5804 application (Figure 3E-H), and they were merely observed after 6 days post-injury (*data not shown*).

### DMX-5804 Promotes MG Proliferation in a YAP-dependent Manner in the NMDA-Injured Retina

To test if the effect of DMX-5804 is YAP dependent, we delivered Cre tagged with enhanced green fluorescence protein (EGFP) into adult Yap^flox/flox^ mouse retina under the control of GFAP promoter by intravitreal injection of adeno-associated viruses (AAVs, type 2/9) (Figure 4A). Loss of YAP staining in SOX9^+^/EGFP^+^ MG was shown in retinal sections, suggesting that the Yap gene in EGFP^+^ MG was successfully removed by Cre recombinase (Figure 4A). In these AAV-treated mice (Figure 4B), we found a decrease of EDU^+^ cells as well as EDU^+^/SOX9^+^ cells in Yap^flox/flox^ mice *vs*. in wild type mice after NMDA and DMX-5804 treatment (Figure 4C-F).

When translocated into the nucleus, YAP binds to the transcription factors TEAD1-4 to promote their activity ^29^. To further determine whether YAP functioned downstream of MAP4K4/6/7 to mediate MG proliferation, we used verteporfin, an inhibitor that prevents YAP-TEAD complex formation, together with DMX-5804 to treat mice for 5 days after NMDA-induced retinal damage (Figure 4G, H); EdU was injected *i.p*. and retinal samples were collected. Compared with the group treated with DMX-5804 only, we observed a decrease in the number of EDU^+^/SOX9^+^ MG in the NMDA-damaged retina in the mice when DMX-5804 and verteporfin were injected together (Figure 3I, J). Taken together, these results confirmed that YAP is the downstream target of DMX-5804 to promote MG proliferation after retinal damage.

### DMX-5804 Induces the Dedifferentiation of MG into a Progenitor-like state in the NMDA-Injured Retina

Pax6 is highly expressed by retinal progenitors in the developing retina and inner retinal neurons (*i.e*., amacrine cells and RGCs) ^30^. The re-emergence of Pax6 expression is a well-documented step during the process of de-differentiation of MG after retinal damage in both fish and chicken *in vivo* ^9,31,32^ as well as in adult rats *in vitro* ^33^. Moreover, unbiased single-cell mRNA sequencing (scRNA-seq) transcriptome analysis has suggested YAP5SA+ MG-derived cells could express Pax6 ^20^. Thus, the expression of Pax6 in MG after DMX-5804 treatment was examined by immunofluorescent staining of retinal sections from NMDA-damaged eyes (Figure 5A). The staining results demonstrated that the YAP^+^/Pax6^+^ cell population expanded in MG with a similar Pax6 fluorescence intensity as the amacrine-rich layer after NMDA/DMX-5804 injections, while much a lower expression level of Pax6 was detected in MG in NMDA/solvent group (Figure 5B, *arrowheads*).

### DMX-5804 Reprograms MG into Retinal Neurons in the NMDA-Injured Retina

To trace the fate of Pax6^+^ MG, Glast-CreERT2^+^/tg;ROSA26R-tdTomato+/tg mice were generated to specifically label MG ^20^ without leakage of neurons (Figure S4A and B). In these mice (Figure 5C), we observed apical translocation of Pax6^+^ MG nuclei to the inner nuclear layer (INL) and outer nuclear layer (ONL) at day 3 after NMDA-induced damage was initiated and DMX-5804 continuous treatment (Figure 5D), which is analogous to a process termed “interkinetic nuclear migration” (INM) during retinal regeneration of zebrafish ^34^. Moreover, we discovered a substantial cohort of tdTomato-positive cells could be labelled by both EdU and Pax6 (Edu^+^/Pax6^+^), suggesting the appearance of proliferating RPC-like cells (Figure 5D). More interestingly, we found some *triply* labeled (EdU^+^/NeuN^+^/tdTomato^+^) cells after 3 days of NMDA/DMX-5804 injections (Figure 5D). Therefore, these results suggested that the conversion of proliferating MG into neurons occurred at an early stage of NMDA/DMX-5804 treatment.

In order to facilitate the observation of cell reprograming process after NMDA or NMDA/DMX-5804 treatment, neuroretinas were flattened and stained to monitor a variety of reprogrammed cells in the whole retina simultaneously (Figure S4C). Consistently, we only discovered reprogrammed tdTomato^+^ MG in the NMDA/DMX-5804 group, which were absent in the NMDA group. In addition,, three types of reprogrammed tdTomato^+^ MG were observed, including EdU^+^/NeuN^−^/tdTomato^+^ MG (Figure S4Da), EdU^+^/NeuN^+^/tdTomato^+^ MG (Figure S4Db), and EdU^−^/NeuN^+^/tdTomato^+^ MG (Figure S4Dc). We suspected these cells could represent three different reprogramming states, respectively: 1) proliferating multipotent MG-derived progenitors; 2) MG-derived neurons after cells finish a cell cycle; and 3) MG-derived neurons without entering a cell cycle. Thus, notably, our results indicated that MG have the ability to transdifferentiate into neurons in both cell cycle-dependent and cell cycle-independent manners after NMDA/DMX-5804 treatment.

### DMX-5804 Induces the Expression of Proneural Transcription Factors in MG in the NMDA-Injured Retina

During development, retinal progenitors express several key proneural transcription factors, including Ascl1, Atoh1/7, Brn3b and Isl1, which are critical to maintain progenitor pool and determine retinal neuron fate ^16,35^. Among them, Ascl1 regulates retinal progenitor pool and induces neuronal fate, while Atoh1/7 has been regarded as a fate determinant of RGC lineage. In addition, Pou4f2 and Isl1 are known to regulate RGC specification and differentiation ^12,16,35^. All of these lineage-specific transcription factors have been reported to affect MG neurogenic capacities previously ^16^.

To test whether MG after NMDA/DMX-5804 treatment acquired the ability to transdifferentiate into neurons, we used droplet digital PCR to determine the expression level of several key transcription factors in isolated MG. Adult Glast-CreERT2+/tg; ROSA26R-tdTomato+/tg mice were induced with tamoxifen followed by NMDA/DMX-5804 injection. Subsequently, tdTomato+ cells were isolated by fluorescence-activated cell sorting (FACS) at different time points (Figure S5A). We detected the increased expression of *Ascl1* from 3 dpi to 10 dpi, and *Atoh7* from 10 dpi to 15 dpi in MG after NMDA/DMX-5804 induction (Figure S5B-D). However, we failed to detect *Atoh1*, *Brn3b* and *Isl1* expression from 0 to 20 dpi (Figure S5B), indicating NMDA/DMX-5804 induced-MG derived neuron-like cells may not reach the maturity. Nonetheless, our data suggested that MG acquired the potential to convert to the RGC lineage after NMDA/DMX-5804 treatment.

### Long-term Tracing of MG Fate after NMDA/DMX-5804 Treatment

To elucidate how efficiently reprogrammed MG could turn into neurons and further determine their eventual fate, we traced MG for 30 days after NMDA/DMX-5804 treatment using Glast-CreERT2^+^/tg;ROSA26R-tdTomato^+^/tg mice (Figure 6A). Although NeuN^+^/tdTomato^+^ cells were absent in the normal (Figure S4B), NMDA-damaged or NMDA/solvent treated retinas (Figure 6B), they appeared in the GCL and INL of retinas after NMDA/DMX-5804 treatment (Figure 6C). We found only a couple of tdTomato^+^ cells co-expressed GAD67, a marker of amacrine cells ^30^ (Figure 6D). In addition, we surprisingly found several tdTomato^+^ cells could be labeled by RNA binding protein with multiple splicing (RBPMS) or βIII-tubulin, two well-acknowledged markers of RGCs ^36,37^, indicating DMX-5804 treated MG might transdifferentiate into RGC-like cells (Figure 6D, *arrowheads*). However, tdTomato^+^/RBPMS^+^ and tdTomato^+^/βIII-tubulin^+^ cells were very rare. Therefore, we set up another paradigm to further confirm the existence of MG-derived RGC-like cells at 30 dpi after NMDA/DMX-5804 treatment. A curtate GFAP promoter was used to increase the label specificity of MG ^10,38^ and MG-specific AAV variants were used (Figure S6A). After intravitreal injection of AAV2/8-short GFAP^−^ EGFP for 3 wk, we detected the strongest green fluorescence signal in the mouse retina (Figure S6B,C). When retinal neurons were stained with NeuN antibodies and MG were stained with Sox9 antibodies, EGFP^+^ signal was only detected in Sox9^+^ cells, further confirming that MG were labeled specifically and their identities were not affected by the injection and treatment with AAVs (Figure S6D). NMDA/DMX-5804 treatment was carried out at the 3^rd^ week after AAVs injection when EGFP signals reached their highest level (data not shown). Subsequently, the retinal samples were collected on 30 dpi (Figure S7A). Immunofluorescent staining of retinal flatmounts or transverse cryosections showed the presence of NeuN^+^/EGFP^+^ cells in the GCL and INL of retinas after NMDA/DMX-5804 treatment, while they were absent after NMDA/solvent treatment (Figure S7B,D,E). Moreover, we also found a small quantity of GAD67^+^/EGFP^+^ cells located in the INL (Figure S7C). Very occasionally, RBPMS^+^/EGFP^+^ cells were observed in the GCL (Figure S7C). Thus, despite a relatively high conversion efficiency from MG into neurons (NeuN^+^ MG-derived cells), only a very small fraction of converted cells could further differentiate into mature neuronal cells expressing the markers of amacrine cells or RGCs.

## DISCUSSION

In zebrafish, reactive MG take three sequential steps to participate in the process of retinal damage repair: (1) acquisition of stem cell properties after retinal damage and proliferation in response to a variety of growth factors and cytokines; (2) amplification of a small population of progenitors through a process called interkinetic nuclear migration in a Pax6-dependent manner; (3) cell cycle exit and neuronal differentiation ^2,39^. Since mammals and zebrafish share the same vertebrate ancestors hundreds of millions of years ago, it is interesting to ask whether mammalian MG retain some hidden regenerative abilities. Here, our results support the claims that MG have the ability to undergo the same process as zebrafish to regenerate neurons, while in the injured adult mouse retina this capability is lost at least partially by YAP inhibition.

The canonical Hippo pathway mediated by MST1/2 has been extensively examined with respect to various biological processes ^40,41^, whereas the physiological and pathological roles of MAP4Ks-mediated YAP phosphorylation remain largely enigmatic. Here, we showed that MAP4K4/6/7, one class of MAP4Ks that regulate YAP activities, controlled the reprogramming of injury-responsive MG in adult mice. Importantly, to our knowledge, this is the first piece of evidence to demonstrate the functions of MAP4Ks-mediated YAP regulation in the mammalian retina. Even more promising, we found that a small molecule inhibitor DMX-5804, which simultaneously inhibits MAP4K4/6/7 ^24^, could effectively promote MG to proliferate and dedifferentiate into RPC-like cells. Moreover, these MG-derived RPC-like cells could spontaneously transdifferentiate into retinal neurons after drug withdrawal.

Previously, transgenic expression of a phospho-deficient variant of YAP (YAP5SA) were reported to stimulate robust proliferation and reprogramming of MG into a progenitor-like state ^19,20^. However, most of the YAP5SA-reprogrammed MG reside in a progenitor-like state and fail to differentiate into neurons due to the constitutively activation of transgenic YAP5SA ^42^. Therefore, one key question remaining after those studies was whether MG are able to transdifferentiate into neurons after they have become reprogrammed into RPC-like cells by regulating the Hippo pathway. Thus, our work has pushed one step forward by demonstrating that mammalian MG retain the intrinsic ability to transdifferentiate into retinal neurons after they enter the RPC-like state. Furthermore, our data support the advantages of chemical induction in comparison with genetic manipulations, especially for cells which have to go through multiple transitional states to regenerate tissues. More importantly, our results provide compelling proof-of-principle to support the feasibility of *in vivo* reprogramming and retinal regeneration by applying a therapeutic intervention based upon use of small molecule chemical cocktails in humans.

RGCs are essential neurons in the retina required for the transmission of encoded visual information from the eye to the brain. However, the inability to regenerate lost or severely degenerated RGCs in mammals, either due to disease or trauma, leads to blindness. In contrast, fish can regenerate their entire retina, including RGCs. Considering the same vertebrate ancestry of fish and mammals, the question of whether RGCs can be regenerated from MG to cure such vision impairments in humans has been a central issue and a quest in the ophthalmic regenerative medicine field many decades. Some studies have suggested that functional RGCs can be regenerated from MG *in vivo* by loss of PTBP1 ^14^ or overexpression Math5:Brn3b ^15^, but the tracing system has raised concerns ^43,44^. Recently, co-expressions of bHLH transcription factors Ascl1:Atoh1 have been shown to regenerate RGC-like cells from MG *in vivo* ^16^. However, it remains unknown whether modulating endogenous signaling pathways by small molecule chemicals can transdifferentiate MG into RGCs. To provide answers for this question, we used a small molecule DMX-5804 to activate YAP and reprogram MG into a progenitor-like state in the first step, and then RPC-like MG exhibited the intrinsic ability to differentiate into retinal neurons after drug withdraw. Droplet digital PCR detected the expression of *Ascl1* and *Atoh7* in these cells (Figure S5). To our surprise, we identified a small subset of MG that transdifferentiated into RGC-like cells, expressing RBPMS and βIII-tubulin (Figure 6D). Based on these findings, we propose that MG indeed retain neuron regenerative abilities, and it is feasible to regenerate retinal neurons in the mammalian retina by using small molecule cocktails, rather than, for example, genetic manipulations.

In addition to DMX-5804, we evaluated another canonical Hippo pathway inhibitor XMU-MP-1, which specifically inhibits MST1/MST2 ^40^. As expected, intraperitoneal administration of XMU-MP-1 suppressed YAP phosphorylation in a dose- and time-dependent manner in adult murine retinas as well (Figure S8A-I). Moreover, the ability of XMU-MP-1 was comparable to DMX-5804 in the ability to promote MG proliferation in NMDA-damaged retinas (Figure S9A-D). However, DMX-5804 showed a higher potency than XMU-MP-1 to promote neuron regeneration in long-term fate tracing experiments (Figure S9E-J). To solve these differences, we found that only DMX-5804, but not XMU-MP-1, modulated JNK pathway activities in adult murine retinas (Figure S10). More specifically, DMX-5804 also suppressed JNK phosphorylation in NMDA-damaged retinas (Figure S11). Previously, it has been reported that inhibition of MAP4K4/6/7 is beneficial for neuroprotection by suppressing apoptosis downstream of the JNK signaling cascade ^45,46^. Accordingly, the levels of JNK phosphorylation in the NMDA-damaged neuroretinas were significantly elevated relative to those in normal (control) retinas during the first 5 days after NMDA injection (Figure S11). Newborn neurons appeared on the 3^rd^ day after NMDA/DMX-5804 administration (Figure 5D; Figure S4C and D). Thus, we suspected that elevated JNK activities made fragile newborn retinal neurons susceptible to cell death in NMDA/XMU-MP-1 treated neuroretinas. Moreover, has been reported that during retinal regeneration, an inflammatory microenvironment affects the survival of newborn neurons in the zebrafish retina ^47^. MAP4Ks play diverse roles in immune cell signaling, immune responses, and inflammation ^48^. Therefore, DMX-5804 treatment may also relieve the inflammation within the microenvironment and protect neurogenesis in the injured retina by inhibiting the MAPK cascades. Taken together, intracellular signaling pathways and extracellular microenvironment might work synergistically to enhance the retinal regenerative efficiency of DMX-5804.

Clinically, a considerable number of patients come to the clinics when they pass the acute phase of retinal injury. Therefore, we were curious about the retinal regenerative capabilities of DMX-5804 when drugs were delivered after the acute phase of retinal damage. It is intriguing to note that DMX-5804 showed higher efficacy than XMU-MP-1 in promoting MG proliferation when both compounds were administrated 2 days post NMDA-induced injury (Figure 13C-F). To interpret this finding, we checked the time course of both canonical and MAP4Ks-mediated Hippo pathway activation after retinal injury, indicated by the phosphorylation level of MOB1 and YAP ^40,41^. The phosphorylation levels of both MOB1 and YAP increased sharply within 24 h after retinal damage (Figure 2B-E; Figure S12A and B). However, MOB1 phosphorylation (pMOB1) was diminished (Figure S12A and B), while YAP phosphorylation remained higher than the normal level 24 h after injury (Figure 2B-E), which might serve as an insurance mechanism to prevent proliferative signals from rebounding in mammals. This inconsistency between the phosphorylation levels of MOB1 and YAP might explain the weaker effect of XMU-MP-1 when both compounds were injected after the acute phase (2 dpi) separately. Thus, these results further support the claims that MAP4Ks-mediated YAP phosphorylation might be a more appropriate target for drug development.

Numerous ways have been invented to stimulate MG de-differentiation and trans-differentiation in mammals to date. However, transgenic tools were used in most of those cases ^10,13,14^. AAVs, which have high affinities for glial cells, have been used as vectors to express pro-proliferation genes or silence endogenous suppressor genes in the damaged retina to assist the de-differentiation, proliferation and trans-differentiation of MG. However, AAVs have only limited cargo capacity to carry foreign genes ^10,49^. As a result, high titers of AAVs expressing various genes or silencing sequences have to be injected intraocularly, since multiple regulatory factors are required to modulate the fate of MG for retinal regeneration, which may also raise biosafety issues ^42,49^. To conquer this problem, the combination of multiple small-molecule chemicals targeting endogenous signaling pathways may be a better choice. Here, we showed that the small-molecule compound DMX-5804 promoted adult mouse MG to reprogram, proliferate and spontaneously trans-differentiate into retinal neurons, with a small fraction expressing the markers of amacrine cells and RGCs after retinal damage. However, the conversion efficiency from MG into mature neurons is relatively low in our hands. Therefore, we propose DMX-5804 might be used as one of several ingredients comprising a cocktail therapy for retinal regeneration. Moreover, the development of appropriate carriers for intraocular delivery might further enhance the efficacy of drug combinations and provide more benefits for clinical practice.

## ACKNOWLEDGMENTS

This work was supported by grants from the National Key R&D Program of China (No. 2018YFA0107304, ZL, No. 2018YFA0107301, WL, No. 2019YFA0111200, YL), National Natural Science Foundation of China (No. 81870627, ZL; No. U20A20363, JH). SJF is the recipient of a Research Career Scientist Award (I K6 BX005787) from the U.S. Department of Veterans Affairs, BLR&D Service. The opinions expressed herein do not reflect those of the Department of Veteran Affairs or the U.S. Government. We thank Dr. Zhen Liu for providing the MIO-M1 cell line. We thank Jingru Huang and Xiang You from Central Lab, School of Medicine, Xiamen University for technical support in confocal imaging. We thank Caiming Wu (State Key Laboratory of Cellular Stress Biology, School of Life Sciences, Xiamen University) for help with droplet digital PCR. We thank Haiping Zheng from Central Lab, School of Medicine, Xiamen University for technical support in fluorescence-activated cell sorting (FACS). We also thank Dr. Wenjun Xiong from Hongkong City University, Dr. Dan Du from Xiamen University, Dr. Kuan Zhang from Army Medical University and Dr. Huaxi Xu from Xiamen University for insightful suggestions and discussions.

## AUTHOR CONTRIBUTIONS

Zuguo Liu, Yi Liao and Houjian Zhang designed projects; Houjian Zhang, Yi Liao, Jiaoyue Hu, Wei Li, and Steven J. Fliesler wrote the manuscript; Houjian Zhang and Yuli Guo performed most of the experiments and data analysis. Yaqiong Yang screened and identified Glast-CreERT2+/tg; ROSA26R-tdTomato+/tg mice for tracing experiment. Jianfeng Wu provided Yap^flox/flox^ mice for YAP knockout experiments. Haiwei Xu provided Glast-CreERT2+/tg mice and useful discussions. Jingbin Zhuang, Yuqian Wang, Jiankai Zhao and Rongrong Zhang performed partial real time PCR, western blot and immunofluorescence.

## Experimental models and subject details

All animals were housed in Xiamen University Laboratory Animal Center; all procedures were approved by the Xiamen University Institutional Animal Care and Use Committee (IACUC) and complied with the *ARVO Statement for the Use of Animals in Ophthalmic and Vision Research*. Glast-CreERT2+/tg;ROSA26R-tdTomato^+^/tg mice were from a mix of Glast-CreERT2^+^/tg C57BL/6J and ROSA26R-tdTomato^+^/tg C57BL/6J mice. Males and females were both used in experiments. All *in vivo* experiments were performed on adult mice that were at least 40 days old.

### MIO-MI cells

MIO-MI cells (RRID: CVCL_0433) were a kind gift from Wenzhou Medical University. The cells were maintained in DMEM containing 10% fetal bovine serum and Penicillin/Streptomycin. For all treatments, 6.0×10^5^ cells per well were seeded into 6-well plates. For siRNA experiments, MIO-M1 cells were transfected using the RNAFit according to the instructions (HANBIO). MAP4K4 and NC siRNA (HH20201222FJCJY-SI01), MAP4K6 and NC siRNA (HH20201222FJCJY-SI02), MAP4K7 and NC siRNA (HH20201222FJCJY-SI03) were synthesized at HANBIO (Shanghai, China). For small-molecule inhibitors treatment, cells were treated with different concentration of DMX-5804 (Cat. No. HY-111754, MedChemExpress). Subsequently, a proper concentration was selected to verify time-dependent effects of DMX-5804.

### AAV production and intravitreal injection

AAV2/9-GFAP-CRE-EGFP and AAV2/9-Syn-Gcamp6s was synthesized at HANBIO (Shanghai, China). AAV2/8-short-GFAP-EGFP was synthesized at OBIO technology (Shanghai, China). Mice were first anesthetized through intraperitoneal injection of pentobarbital sodium (1% in PBS). Tropicamide (0.5%) was then topically administered for pupillary dilation. While applying gentle pressure around the eye socket to extrude the eye, a 33-gauge needle was passed through the sclera just behind the limbus, into the vitreous cavity. Injection of 2 μl of AAV was made with direct observation of the needle in the center of the vitreous cavity. For NMDA intravitreal injection, injection of 2 μl of NMDA (100 mM, Cat. No. M3262-25mg, Sigma Aldrich) was made in the center of the vitreous cavity.

## Method details

### The Compounds preparation and intraperitoneal injection

DMX-5804, XMU-MP-1 and verteporfin were dissolved in solvent with the order: 10% DMSO, 40% PEG300, 5% Tween-80 and 45% PBS for mother solution. DMX-5804, XMU-MP-1 and the solvent control were each administered by intraperitoneal injection, as 2 mg/kg dose, 6 h apart for different consecutive days according to experimental designs, while verteporfin and the solvent control were each administered by intraperitoneal injection, as 2 mg/kg dose, 6 h apart but with 1 hour interval before DMX-5804 injection. To label cells in S phase, 5-ethynyl-2’-deoxyuridine (EdU) (APExBIO, Cat. No. B8337) was injected intraperitoneally at a concentration of 50 mg/kg body weight and mice were euthanized by cervical dislocation 12 or 24 h later.

### RNA extraction and RT-qPCR

Total RNA was extracted from mouse neural retina using RNeasy mini kit (QIAGEN). Total RNA was reverse transcribed in the presence of PrimeScript™ RT reagent Kit with gDNA Eraser (Perfect Real Time) (Takara, Cat. No. RR047B). For each RT-qPCR reaction, 1.5 ng of cDNA was used in triplicates in the presence of SYBR Green qPCR Mix (Bio-Rad) on an Applied Biosystems Real-Time PCR Detection System (Thermo Fisher Scientific). The primers used to detect mRNA expression levels were MAP4K4 (Forward: 5’-GAAGGGTGCGTGCACTATAAA-3’; Reverse: 5’-TCCACAGTGAGATCCACCAG-3’), MAP4K6 (Forward: 5’-TCCATGCTGTGGATGTTGAT-3’; Reverse:5’-CGCCCGTAAGTGTTGACATA-3’), MAP4K7 (Forward:5’-GCGAGGAAGAGGAGGAAGAT-3’; Reverse:5’-TGAAACTGTCCGCATGAGAG-3’). 18sRNA (Forward:5’-TCCATGCTGTGGATGTTGAT-3’; Reverse:5’-CGCCCGTAAGTGTTGACATA-3’) were used for normalization. Differential expression analysis was performed using the △△CT method. Relative expression of each gene in each sample was calculated using the mean of the controls as the reference. RT-qPCR experiments were performed on at least 3 mice per condition, allowing for statistical analysis.

### Western blotting

For cells, the culture medium was removed and cells were briefly washed in PBS containing protease (Thermo Fisher Scientific, Cat. No.1862209) and phosphatase inhibitors (Thermo Fisher Scientific, Cat. No.78420). Cells were lysed using a cold lysis buffer (Thermo Fisher Scientific, Cat. No.1861603) containing protease and phosphatase inhibitors and the cells were flash-frozen and kept at −80 °C until used.

For retinas, neural retinas were dissected in PBS and briefly washed in PBS containing protease and phosphatase inhibitors. Samples were put into ice-cold lysis buffer containing protease and phosphatase inhibitors and the retinas were flash-frozen (liquid nitrogen) and then stored at −80 °C until used.

For protein extraction, samples were thawed on ice for 10 min in lysis buffer containing protease and phosphatase inhibitors, then cells or retinas were homogenized and centrifuged at 4 °C (2000 rpm, 10 min), and the supernatants were transferred to a new tube. Clarified protein lysates were quantified by the Bradford assay.

For Western blotting, 20 mg protein was loaded onto 8-10% SDS-PAGE gels for electrophoresis and subsequently transferred (90 V, 2-3 h) onto PVDF western blotting membranes (Roche, Cat. No. 3010040001) using the Criterion™ System BioRad. Membranes were washed in TBST pH 7.4 (20 mM Tris, 137 mM NaCl, 0.1% Tween 20) and blocked for 1 h at room temperature (RT) with 5% non-fat milk in TBST to reduce non-specific antibody binding. At least 3 independent controls and treated samples were probed with the following primary antibodies (in 5% milk, overnight at 4°C): YAP (1:500, Cell signaling Tec., Cat. No. 14074, RRID: AB_2650491), p YAP (1:2000, Cell signaling Tec., Cat. No. 13008, RRID: AB_2650553), MAP4K4 (1:500, Cell Signaling Tec., Cat. No. 3485, RRID: AB_2140972), MAP4K6 (1:500, Novus, Cat. No. NBP1-22989, RRID: AB_1726507) and MAP4K7 (1:500, Genetex, Cat. No. GTX13141, RRID: AB_383061), MOB1 (1:1000; Cell signaling Tec., Cat. No. 13730, RRID: AB_2783010), p MOB1 (1:1000, Cell signaling Tec., Cat. No. 8699, RRID: AB_11139998). Vinculin (1:5000, Cell Signaling Tec., Cat. No. 13901, RRID: AB_2728768) was used as loading control. After primary antibody incubation, membranes were washed in TBS-T, incubated with HRP-conjugated secondary antibodies (1-h, RT), detected with Clarity™ Western ECL Substrate (Bio-Rad, Cat. No. 170-5060) and imaged with the Chemidoc Touch Imager (Bio-Rad Laboratories Inc, Hercules, CA, USA).

### Immunofluorescence staining on cells

MIO-M1 cells seeded in 6-well culture plates were fixed in 4% paraformaldehyde (PFA) in PBS for 10 min. The PFA was removed, and the plates were washed with PBS. Fixed cells were treated with blocking buffer containing 2% BSA in PBS for 1 h on ice. Plates were incubated for 24 h in primary antibodies in a humidified chamber. The following antibodies and concentrations were used: Glutamine Synthetase (1:500, Millipore, Cat. No. MABN1182, RRID: AB_2110656), GFAP (1:500, Millipore, Cat. No. G6171, RRID:AB_1840893), MAP4K4 (1:500; Cell Signaling Tec., Cat. No. 3485, RRID: AB_2140972), MINK1 (1:500; Novus, Cat. No. NBP1-22989, RRID: AB_1726507) and TNIK (1:500; Genetex, Cat. No. GTX13141, RRID: AB_383061), YAP (1:100, Santa Cruz, Cat. No. 101199, RRID: AB_1131430). After washing with PBS, anti-mouse (488, 1:300, Thermo Fisher Scientific, Cat. No. A21022, RRID: AB_141607) or anti-rabbit (488, 1:300, Thermo Fisher Scientific, Cat. No. A11008, RRID: AB_143165 or 594, 1:300, Thermo Fisher Scientific, Cat. No. A11012, RRID: AB_141359) secondary antibodies were applied, and signals were visualized using a Zeiss LSM 880+Airyscan.

### Immunofluorescence staining on tissue sections

For the paraffin sections, enucleated eyes were immersed in 4% paraformaldehyde (PFA) in 0.1 M PBS overnight at room temperature (RT), corneas and lenses were removed. After fixation, eyecups were administered tissue dehydration in accordance to the order: 70% ethyl alcohol (2 h), 80% ethyl alcohol (1 h), 95% ethyl alcohol (1 h), 100% ethyl alcohol (1 h, with 3 repeats), xylene (30 min, with 3 repeats), dehydrated samples were embedded with paraffin at 60 °C (1 h, with 3 repeats), and then mounted in paraffin tissue molds and stored at RT before sectioning. Paraffin sections were cut at 8 μm thickness on a microtome and mounted on Superfrost Plus slides (CITOTEST, Cat. No. 188105). Before immunofluorescence, slides were dewaxed in accordance to the order: xylene (15 min, 2 repeats), 100% ethyl alcohol (10 min), 95% ethyl alcohol (5 min), 80% ethyl alcohol (5 min), 70% ethyl alcohol (5 min) and antigen-retrieval was performed in antigen retrieval buffer (3 g sodium citrate and 0.4 g citric acid in 1 L deionized water) at 100 °C for 20 min. For EdU, sections were permeabilized for 30 min in 0.5% Titron-X100 diluted in 0.1 M PBS, and then washed using 3%BSA diluted in 0.1 M PBS. The Click-iT EdU Alexa Fluor® 594 (Thermo Fisher Scientific, Cat. No.C10639) was used following the manufacturer’s instructions.

For the cryosections, enucleated eyes were immersed in 4% PFA in 0.1 M PBS for 1 h at RT, corneas and lenses were removed. After fixation, eyecups were washed in PBS 3 times for 10 min on ice, and then cryoprotected by immersing in 30% sucrose until the tissues sank to the bottom of the tubes. Then, tissues were embedded in O.C.T. for 15-20 min and then flash-frozen in liquid nitrogen and stored at −80 °C before sectioning. Cryosections were cut at 20 μm thickness on a cryostat and mounted on Superfrost Plus slides (CITOTEST, Cat. No. 188105).

For immunofluorescence, sections were rinsed with PBS and incubated for 2 h at 4°C with 2% bovine serum albumin (BSA), and 0.5% Triton X-100 diluted in 0.1 M PBS (pH 7.4). Slides were incubated for 24 h in primary antibodies in a humid chamber. The following antibodies and concentrations were used: YAP (1:100, Santa Cruz, Cat. No. 101199, RRID: AB_1131430), SOX9 (1:500, Abcam, Cat. No. ab185966, RRID: AB_2728660), GFAP (1:500, Millipore, Cat. No. G6171, RRID:AB_1840893), NeuN (1:500, Abcam, Cat. No. ab177487, RRID: AB_2532109), GAD67 (1:100, Abcam, Cat. No. ab26116, RRID: AB_448990), RBPMS (1:500, Proteintech, Cat. No. 15187-1-AP, RRID: AB_2238431). Next, slides were rinsed 3 times in PBS and blocked for 30 min in 2% BSA, and 0.5% Triton X-100 diluted in 0.1 M PBS (pH 7.4) before incubation in secondary antibodies. Anti-mouse (488, 1:300, Thermo Fisher Scientific, Cat. No. A21022, RRID: AB_141607 or 594, 1:300, Thermo Fisher Scientific, Cat. No. A11032, RRID: AB_2534091) or anti-rabbit (Alexa Fluor-488, 1:300, Thermo Fisher Scientific, Cat. No. A11008, RRID: AB_143165 or Alex Fluor-594, 1:300, Thermo Fisher Scientific, Cat. No. A11012, RRID: AB_141359) secondary antibodies were applied and incubated for 1 h in the dark at RT. Hoechst 33342 (1:2000, Thermo Fisher Scientific, Cat. No. H3570) was included in the secondary antibody solution at a 1:2000 concentration. After incubation with secondary antibodies, slides were rinsed 3 times in PBS. Slides were coverslipped with VECTASHIELD® antifade mounting medium (Vector Labs, Cat. No. H-1000-10). Retinal sections from controls and treated mice were immunolabeled in parallel to insure identical processing.

### EdU staining and immunofluorescence on whole mount

Immunostaining on whole mount was performed using standard procedures with the following modifications: (i) enucleated eyes were immersed in 4% PFA in 0.1 M PBS for 1 h at RT, then retinas were isolated from the eyecup, and incised radially into four radial pieces; (ii) the retinas were first incubated overnight in a blocking solution containing 2% BSA and 0.5% Triton X-100 in PBS (pH 7.4); (iii) For EdU, retinas were washed using 3% BSA diluted in 0.1 M PBS. The Click-iT EdU Alexa 488 (Thermo Fisher Scientific, Cat. No.C10638), Click-iT EdU Alexa 594 (Thermo Fisher Scientific, Cat. No.C10639), or Click-iT EdU Alexa 647 (Thermo Fisher Scientific, Cat. No.C10640) was used following the manufacturer’s instructions. (iii) The retinas then were incubated in a mixture of primary antibodies. The following antibodies and concentrations were used: SOX9 (1:500, Abcam, Cat. No. ab185966, RRID: AB_2728660), NeuN (1:500, Abcam, Cat. No. ab177487, RRID: AB_2532109), GAD67 (1:100, Abcam, Cat. No. ab26116, RRID: AB_448990), RBPMS (1:500, Proteintech, Cat. No. 15187-1-AP, RRID: AB_2238431). Nuclei were counterstained with 1:2000 Hoechst (1:2000, Thermo Fisher Scientific, Cat. No. H3570). Next, retinas were rinsed 3 times in PBS and blocked for 30 min in 2% BSA, and 0.5% Triton X-100 diluted in 0.1 M PBS (pH 7.4) before incubation in secondary antibodies. 1:300 dilutions of Alexa Fluor 488 or 594 conjugated secondary antibodies (antibodies were from Thermo Fisher Scientific which has been described above) were applied and incubated for 1 h in the dark at room temperature. Hoechst was included in the secondary antibody solution at a 1:2000 dilution. Finally, Slides were coverslipped with VECTASHIELD® antifade mounting medium (Vector Labs, Cat. No. H-1000-10) on Superfrost Plus slides (CITOTEST, Cat. No. 188105) before observation.

### Fluorescence-activated cell sorting (FACS)

Following euthanasia, retinas were dissociated with Papain/DNase for 30 minutes at 37°C. The tissue was then gently triturated for adequate dissociation before Ovomucoid was added. Suspending cells were spun down at 300g and then resuspended in Neurobasal/Ovomucoid/ DNase solution and passed through a 35 microns filter. FACS was then performed on a MoFlo Astrios EQS (Beckman).

### RNA extraction and droplet digital PCR

The RNeasy Plus Micro Kit (cat. no. 74034, QIAGEN) was used for RNA extraction from isolated MG. Total mRNA was reverse transcribed in the presence of PrimeScript™ RT reagent Kit with gDNA Eraser (Perfect Real Time) (Takara, Cat. No. RR047B). Droplet digital PCR was used to measure the levels of transcription factors in isolated MG following a previous method (Fisher et al., 2015). The primers used to amplify the cDNA sequences were *Ascl1* (Forward: 5′-GAACTGATGCGCTGCAAAC-3′; Reverse 5′-CGTCTCCACCTTGCTCATCT-3′); *Atoh1* (Forward: 5′-CCACAGCTTCCTGCAAAAAT-3′; Reverse: 5′-GAGTAACCCCCAGAGGAAGC-3′): *Atoh7* (Forward: 5′-AGTGGGGCCAGGACAAGA-3′; Reverse: 5′-GGGTCTACCTGGAGCCTAGC-3′); *Brn3b* (Forward: 5′-CAGCAGTTCCAGCAGCAGT-3′; Reverse: 5′-ATGGTGGTGGTGGCTCTTAC-3′); *Isl1* (Forward: 5′-AGCTGGAGACCCTCTCAGTC-3′; Reverse: 5′-TGCTTCTCGTTGAGCACAGT-3′). PCR was run using PerfeCta Multiplex qPCR ToughMix (Quanta Biosciences, Gaithersburg, MD, USA). Droplet digital PCR was performed on a Naica Crystal Digital PCR system™ (Stilla Technonologies, Villejuif, France). Stilla’s Crystal Miner® software was used to measure the droplet identification and fluorescence intensity.

### Imaging

Confocal images were acquired using a Zeiss LSM 880+Airyscan. For standardization between samples, all images were acquired with the same laser power, detector gain, scan speed, and pinhole size. Z stack images with an identical step-size and thickness were acquired for each retinal section and a maximal intensity projection were obtained.

### Quantification and statistical analysis

For all figure panels, images were imported into Adobe Photoshop CS software for identical and minimal processing. For each control, mutant, or treated experimental group, three to six independent retinas from different mice were examined and the results presented in the manuscript are representative images from at least three separate experiments. A Student’s *t*-test was used to compare measurements between controls and mutant mice. *P* values < 0.05 were considered as statistically significant.

Quantifications of the number of labeled cells in stretched preparation of mouse retinas were calculated from 5 different fields of per retina and using 3 retinas per condition.

For quantification of nuclear YAP immunofluorescence, ZEN2.3 lite software was used to quantify YAP pixel intensities when coincident with SOX9^+^ pixels (to denote nuclear localization). For EdU quantification of confocal images, EdU^+^ cells were manually counted in each image and retinal area was measured in ZEN2.3 lite software to determine EdU^+^ cells per mm^2^. To measure pixel co-localization across MG within a given retina ZEN2.3 lite software was used to plot pixel intensities along a line drawn across the image and Pearson’s Correlation Coefficient (R) was calculated. For quantification of relative pixel intensity per mm^2^, total pixel intensity was measured in ZEN2.3 lite software and divided by the total area of each image.

For quantification of western blotting, images were taken with the Chemidoc Touch Imager (Bio-Rad Laboratories Inc, Hercules, CA, USA) and scanned at high-resolution for protein band size and signal intensity. Densitometry measurements were obtained using the Image LabTM software (Bio-Rad Laboratories Inc, Hercules, CA, USA). All bands were normalized to their corresponding loading controls. The results shown are from at least 3 biological replicates. The Student’s *t*-test was used to determine differences between mutants and controls; the threshold for statistical significance was set as *p*-values < 0.05. For quantification of mRNA by qPCR, each q-RT-PCR reaction was performed in triplicate for each independent control (n=3) or treatment group (N = 3) cDNA samples. The mean ^ΔΔ^ CT values were normalized against the housekeeping genes 18s rRNA and corresponding ^ΔΔ^CT values were log_2_-transformed to obtain fold change values. For data analysis, the Student’s *t*-test was used to determine relative gene expression ratios and a *p*-value of < 0.05 was considered as statistically significant.

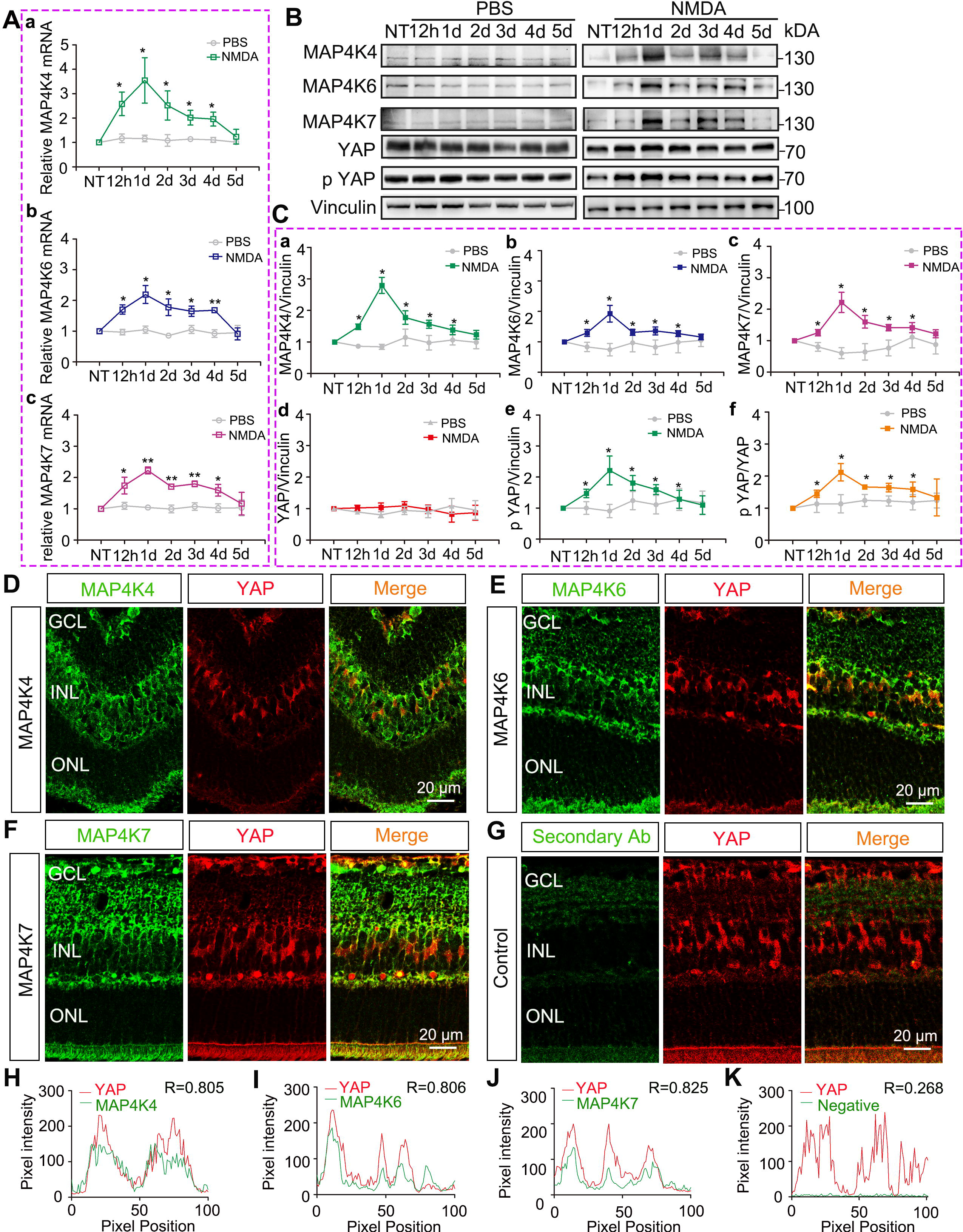

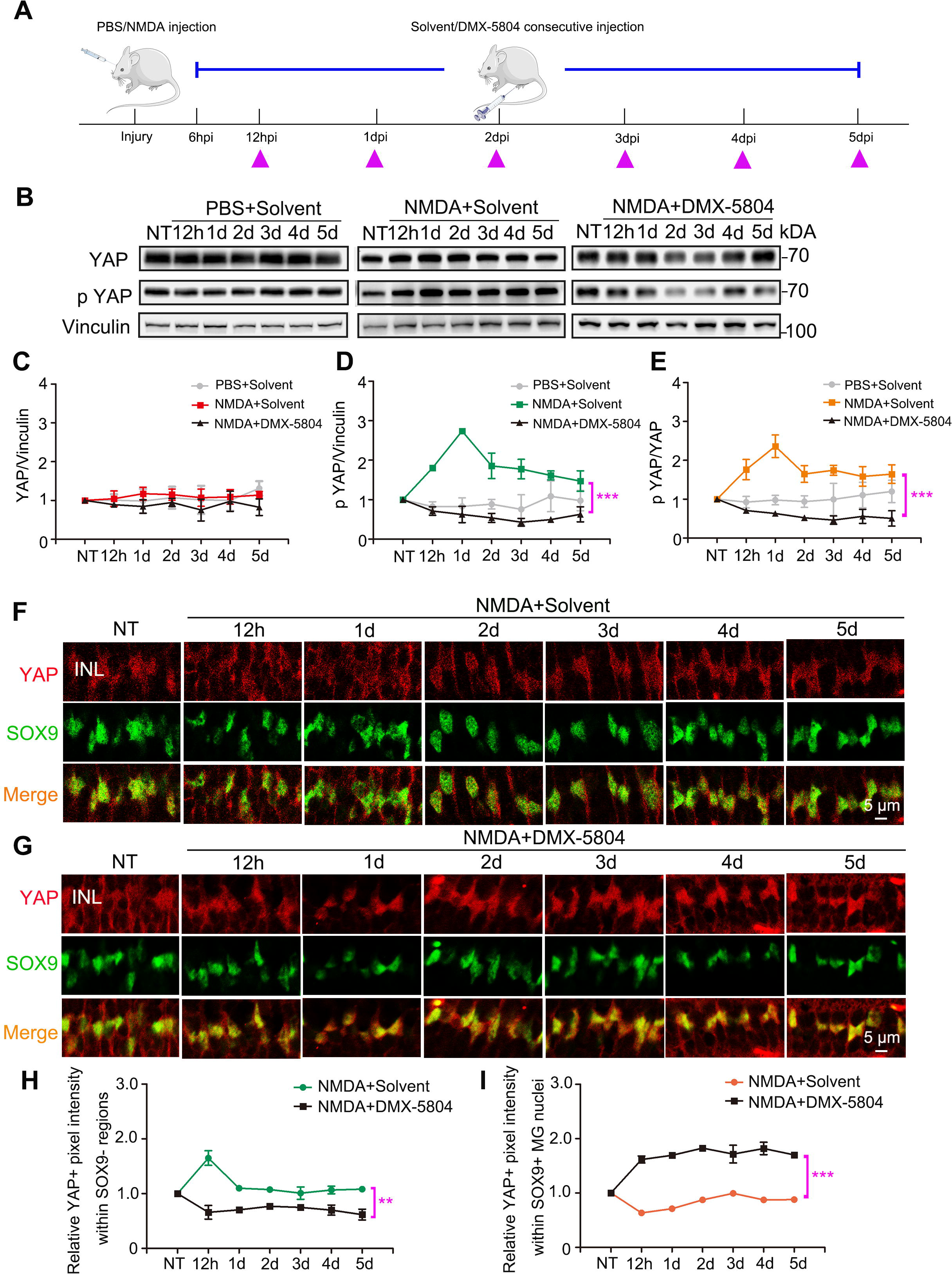

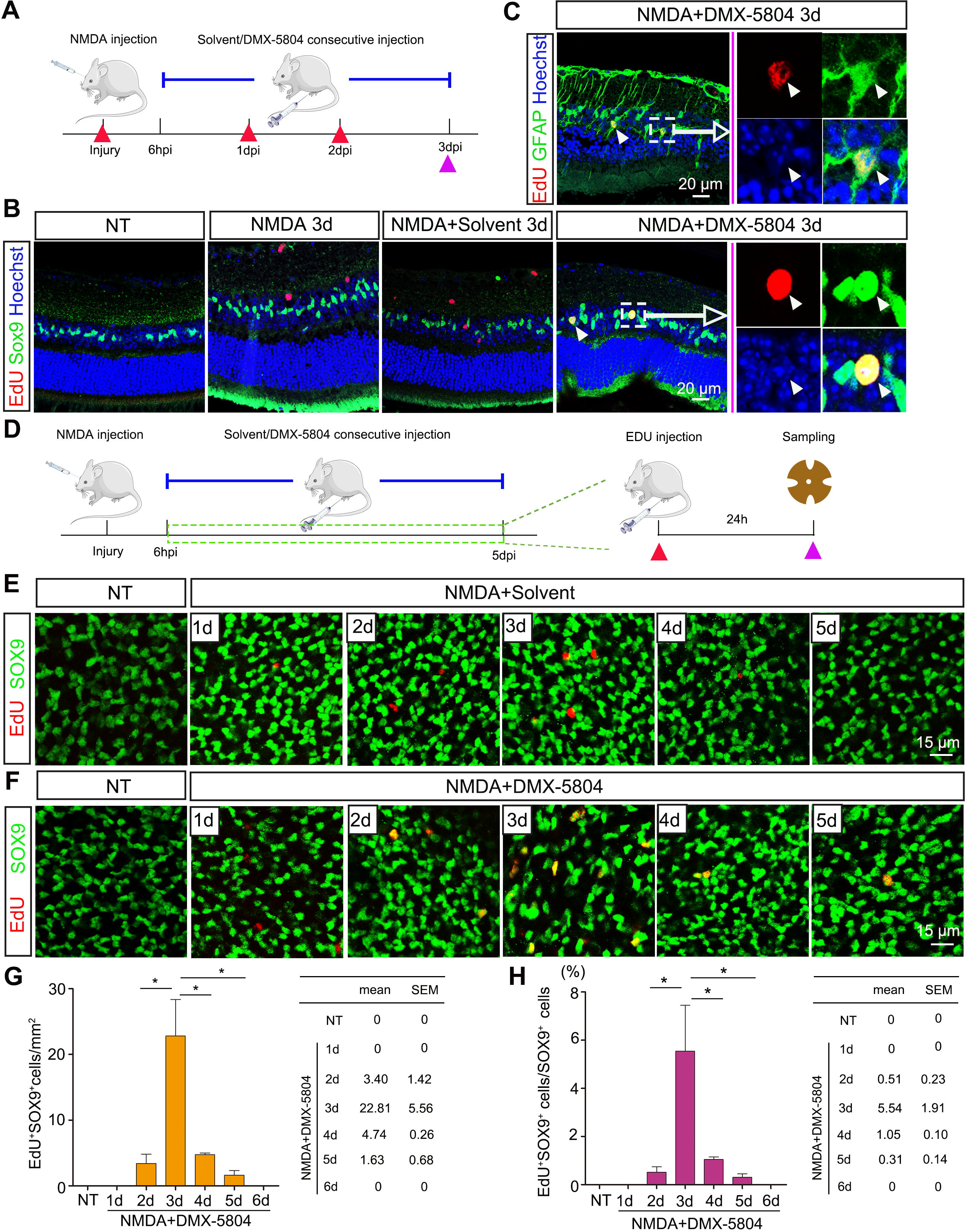

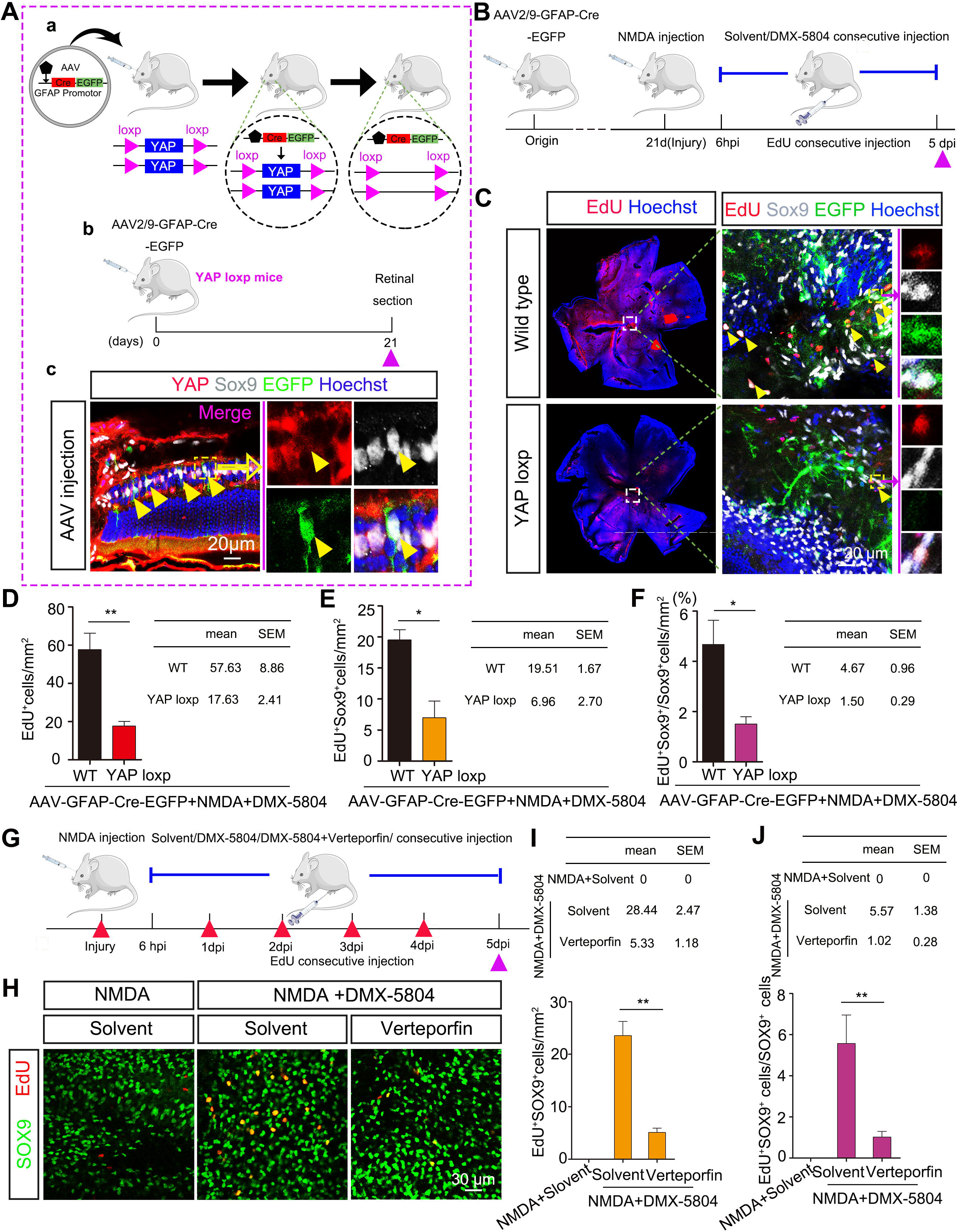

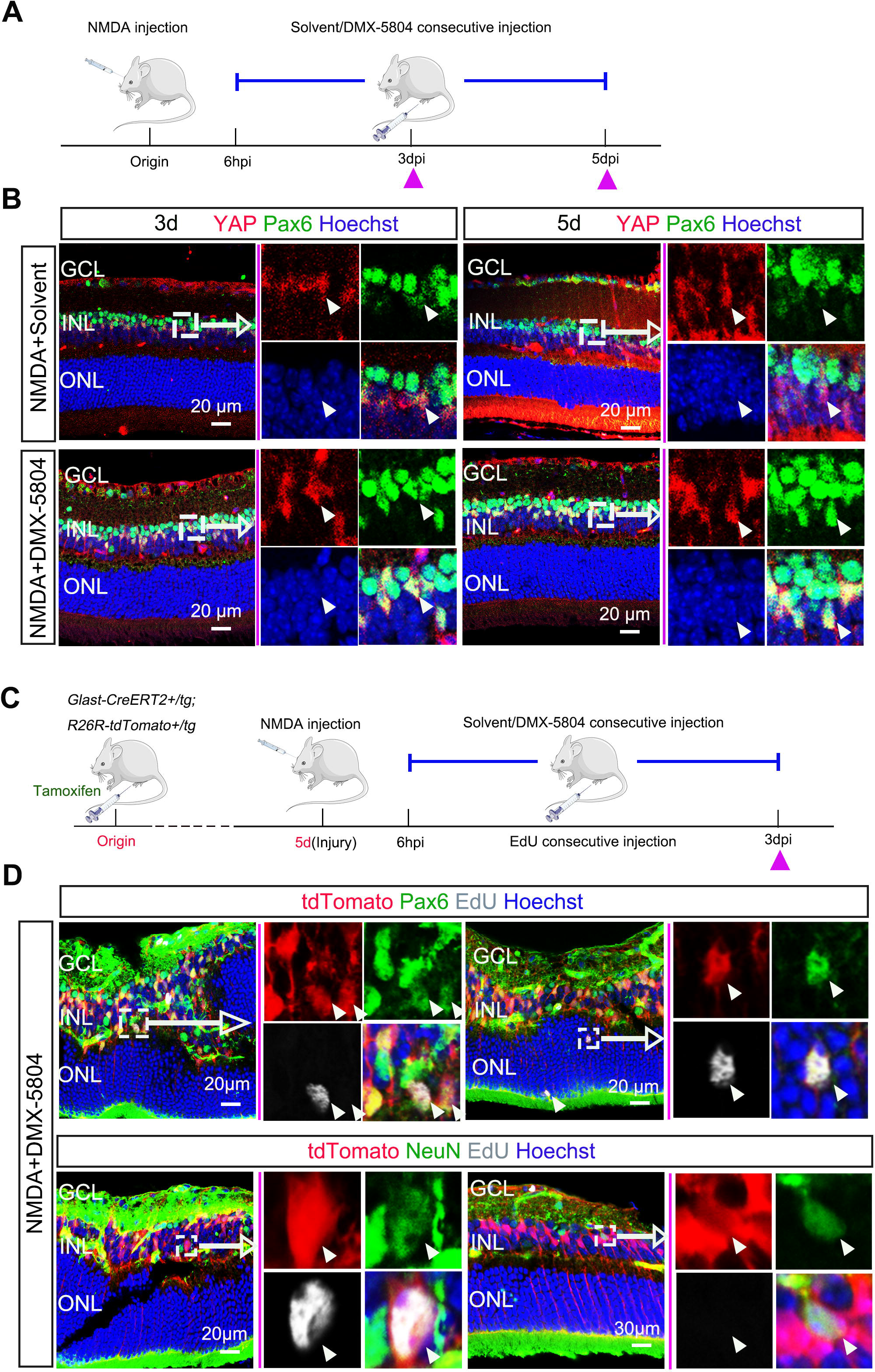

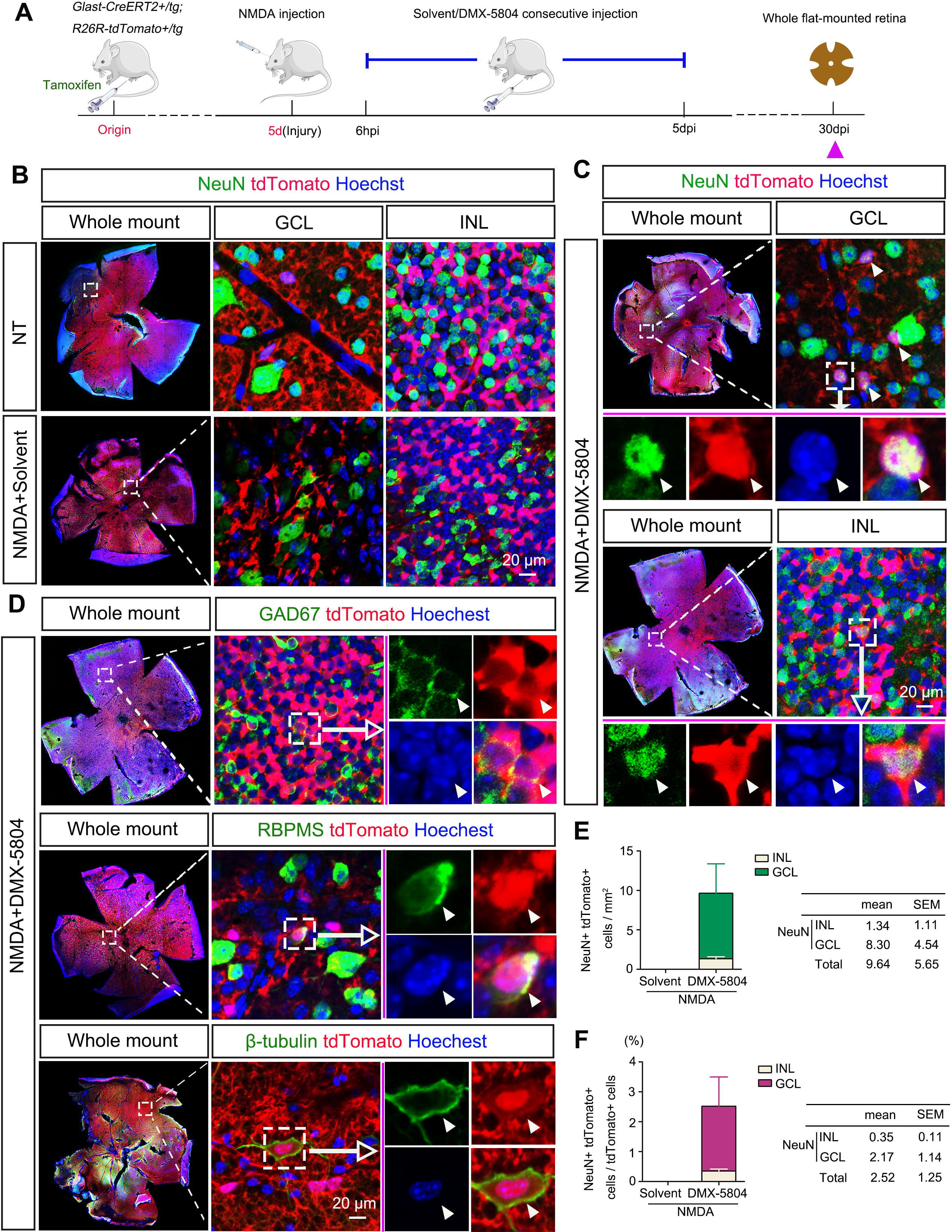

