## Supplementary material for "MAP4Ks inhibition promotes retinal neuron regeneration from Müller glia in adult mice": Figure legend

**Figure 1.** **MAP4K4/6/7 expression and YAP phosphorylation after retinal injury in MG.**

(**A**) Time course analysis of MAP4K4(a)/6(b)/7(c) mRNA in the NMDA-injured retina.

(**B**) Western blots analysis of MAP4K4/6/7, p YAP and YAP in the NMDA-injured retina.

(**C**) Quantification of MAP4K4(a)/6(b)/7(c), p YAP and YAP protein expression(d-f) in (**B**).

(**D-G**) Immunofluorescence staining of MAP4K4 (**D**), MAP4K6 (**E**), MAP4K7 (**F**) together with YAP in the sections of murine retina. MG were specifically stained with YAP. The section stained with secondary antibody (**G**) was served as negative control.

(**H-K**) Co-localization analysis of MAP4K4/6/7/negative control and YAP in (**D-G**).

For qRT-PCR, mRNA levels were compared with the no-treatment (NT) control and depicted as fold-change ± SEM (n = 3 independent pools of samples per group; Student’s *t* test). For Western blots, levels were given as a.u. ± SEM in comparison with PBS or no-treatment (NT) controls (n = 3 independent pools of samples per group; Student’s *t-*test). For co-localization analysis, evaluation results were given as Pearson correlation coefficient. **p*≤0.05, ***p*≤0.01, ***p≤0.001.

**Figure 2. The translocation of YAP into the nuclei of MG after NMDA/DMX-5804 treatment.**

(**A**) Timeline diagram of the experimental procedure used in (**B-I**). Wildtype mice were intravitreally injected with NMDA. Solvent control or DMX-5804 was injected every 6 hours intraperitoneally from 6 h post-NMDA injection (hpi) until 5 days post-NMDA injection (dpi). The purple triangles represent the time points of the sampling.

(**B**) Western blot analysis of p YAP and YAP in the murine neuroretina treated with PBS/solvent, NMDA/solvent and NMDA/DMX-5804.

(**C-E**) Quantification of YAP, p YAP (normalized to vinculin) and p YAP/YAP ratio in (**B**).

(**F**) Immunofluorescence staining of YAP (*red*; YAP+) and SOX9 (*gree*n; SOX9+) at indicated time points in the NMDA-injured murine retina.

(**G**) Immunofluorescence staining of YAP (*red*; YAP+) and SOX9 (*gree*n; SOX9+) at indicated time points in the NMDA-injured murine retina treated with DMX-5804.

(**H**) Quantification of relative YAP+ pixel intensity in SOX9- MG nuclei in (**F**) and (**G**).

(**I**) Quantification of relative YAP+ pixel intensity in SOX9+ MG nuclei in (**F**) and (**G**).

For western blots, levels were given as a.u. ± SEM in comparison with solvent or no-treatment (NT) (n = 3 independent pools of samples per group; one-way ANOVA test). For pixel intensity measurements, levels were given as mean ± SEM (n = 3 per group; one-way ANOVA test). The one-way ANOVA test was performed between the NMDA+Solvent group and the NMDA+DMX-5804 group. **p*≤0.05, ***p*≤0.01, ****p*≤0.001.

**Figure 3.** **Proliferation of MG in the NMDA-injured murine retina with or without DMX-5804 treatment.**

(**A**) Timeline diagram of the experimental procedures used in (**B, C**). Wildtype mice were intravitreally injected with NMDA. Solvent control or DMX-5804 was injected following the same protocol depicted in Figure 2(**A**). EdU was injected intraperitoneally every 24 h after NMDA injury till mice scarification. The *red* triangles represent the time points of the EdU injection and the *purple* triangles represent the time points of the sampling.

(**B**) EdU labeling (*red*) and SOX9 immunofluorescence (*green*) on retinal sections after NMDA, NMDA/solvent or NMDA/DMX-5804 treatment.

(**C**) EdU labeling (*red*) and SOX9 immunofluorescence (*green*) on retinal sections after NMDA/DMX-5804 treatment.

(**D**) Timeline diagram of the experimental procedures used in (**E-H**). Wildtype mice were intravitreally injected with NMDA. Solvent or DMX-5804 was injected using the same method depicted in Figure 2(**A**). A single dose of EdU was injected intraperitoneally 24 h before mice were sacrificed.

(**E**) EdU labeling (*red*) and SOX9 immunofluorescence (*green*) on whole flat-mounted retinas at indicated time points after NMDA/solvent treatment.

(**F**) EdU labeling (*red*) and SOX9 immunofluorescence (*green*) on whole flat-mounted retinas at indicated time points after NMDA/DMX-5804 treatment.

(**G**) Quantification of the number of cells positive for both EdU labeling (EdU+; *red*) and SOX9 immunofluorescence (SOX9+; *green*) per mm^2^ in (**E**).

(**H**) Quantification of the percentage of cells positive for both EdU labeling (EdU+; *red*) and SOX9 immunofluorescence (SOX9+; *green*) per mm^2^ in (**F**).

For quantification of EdU+ SOX9+ cells, levels were given as mean ± SEM (n = 3 retinas per group; Student’s *t*-test). **p*≤0.05, ***p*≤0.01, ****p*≤0.001.

**Figure 4. NMDA/DMX-5804 induced MG proliferation is absent in YAP conditional knockout mice.**

(**A**) Conditional YAP knockout strategies and representative results. Schematic diagram showing conditional YAP knockout strategies and tracing system in mouse Müller glia using AAVs(a,b). YAP (*red*), Sox9 (*gray*) and Cre (EGFP) immunofluorescence in mouse retinal sections after pAAV-short GFAP-MCS-EGFP-3FLAG injection(c). The *red* triangles represent the time points of the EdU injection and the *purple* triangles represent the time points of the sampling.

(**B**) Timeline diagram of the experimental procedures used in (**D-G**). Wild type or YAP*^flox/flox^* mice were intravitreally injected with pAAV-GFAP-Cre-T2A-EGFP. NMDA was intraocularly administered 3 wk later. Then DMX-5804 was injected every 6 h intraperitoneally from 6 hpi to 5 dpi after NMDA injection. After that, mice were sacrificed and the retinas were analyzed.

(**C**) EdU labeling (EdU+; *red*) and SOX9 immunofluorescence (SOX9+; *gray*) on whole flat-mounted retinas infected with pAAV-GFAP-Cre-T2A-EGFP after NMDA injury and DMX-5804 treatment.

(**D**) Quantification of the number of cells positive for EdU labeling (EdU+; *red*) per mm^2^ in

(**E**) Quantification of the number of cells positive for both EdU labeling (EdU+; *red*) and SOX9 immunofluorescence (SOX9+; *gray*) per mm^2^ in (**B**).

(**F**) Quantification of the percentage of EdU+ SOX9+ cells in SOX9+ cells per mm^2^ in (**B**).

(**G**) Timeline diagram of the experimental procedures used in (**G-J**). Wild type mice were intravitreally injected with NMDA. DMX-5804/solvent or DMX-5804/verteporfin were delivered every 6 h intraperitoneally from 6 hpi to 5 dpi. EdU was injected intraperitoneally every 24 h until 5 days after NMDA injury.

(**H**) EdU labeling (EdU+; *red*) and SOX9 immunofluorescence (SOX9+; *green*) on whole flat-mounted retinas 5 days after NMDA injection and DMX-5804/solvent or DMX-5804/verteporfin treatment.

(**I**) Quantification of the number of cells positive for both EdU labeling (EdU+; *re*d) and SOX9 immunofluorescence (SOX9+; *green*) per mm^2^ in (G).

(**J**) Quantification of the percentage of EdU+ SOX9+ cells in SOX9+ cells per mm^2^ in (**G**).

For quantification of EdU+ SOX9+ cells, levels were given as mean ± SEM (n = 3 retinas per group; Student’s *t*-test). **p*≤0.05, ***p*≤0.01, ****p*≤0.001.

**Figure 5.** **MGs enter into retinal progenitor cell-like state and transdifferentiate into neurons after NMDA/DMX-5804 treatment.**

(**A**) Timeline diagram of the experimental procedure used in (**B**). The mice were treated following the experimental procedures depicted in Figure 2(**A**). The *purple* triangles represent the time points of the sampling.

(**B**) YAP (*red*) and Pax6 (*green*) immunofluorescence in the NMDA-injured retinas after solvent or DMX-5804 treatment.

(**C**) Timeline diagram of the experimental procedures used in (**D**). Glast-CreERT2+/tg;ROSA26R-tdTomato+/tg mice were intraperitoneally injected with Tamoxifen for 5 consecutive days. Then NMDA was intraocularly administered. DMX-5804 was injected every 6 h intraperitoneally from 6 hpi until 3 dpi after NMDA injection. The *purple* triangles represent the time points of the sampling.

(**D**) EdU labeling (*gray*) and Pax6 (*green*)/NeuN (*green*) immunofluorescence in the NMDA-injured retinas after 3 days of DMX-5804 treatment.

**Figure 6.** **Long-term tracing of MG Fate after NMDA/DMX-5804 Treatment.**

(**A**) Timeline diagram of the experimental procedures used in (**B-D**). Glast-CreERT2+/tg;ROSA26R-tdTomato+/tg mice were intraperitoneally injected with Tamoxifen for 5 consecutive days. Then NMDA was intraocularly administered. DMX-5804 was injected every 6 h intraperitoneally from 6 hpi until 5 dpi after NMDA injection. Thirty (30) days after NMDA injection, mice were sacrificed and retinal samples were analyzed. The *purple* triangles represent the time points of the sampling.

(**B,C**) NeuN (*green*) immunofluorescence on whole flat-mounted NMDA-injured retinas after Solvent or DMX-5804 treatment.

(**D**) GABAergic cell marker GAD67 (*green*) or retinal ganglion cell marker (RBPMS/βⅢ-tubulin, *green*) immunofluorescence on whole flat-mounted NMDA-injured retinas after DMX-5804 treatment.

(**D**) Quantification of the number of NeuN+tdTomato+, GAD67+tdTomato+ cells or RBPMS+/β-tubulin+tdTomato+ cells per mm^2^ in (**B-D**).

(**E**) Quantification of the percentage of NeuN+tdTomato+, GAD67+tdTomato+ cells or RBPMS+/β-tubulin+tdTomato+ cells in tdTomato+ cells per mm^2^ in (**B-D**).

For quantification of NeuN+ tdTomato+ cells, levels were given as mean ± SEM (n = 3 retinas per group).

**SUPPLEMENTAL FIGURES**

**Figure S1. MAP4K4/6/7 regulates YAP phosphorylation in MIO-M1.**

(**A**) Immunofluorescence staining of Müller glia marker GS, GFAP, as well as MAP4K4/6/7 in MIO-M1.

(**B-D**) Western blot analysis of p YAP and YAP (**B**) and quantification (**C**; normalized to vinculin) in MIO-M1 treated with MAP4K4 siRNAs.

(**E, F**) Western blot analysis of p YAP and YAP (**E**) and quantification (**F**; normalized to vinculin) in MIO-M1 treated with MAP4K4/6/7 siRNAs.

(**G**) YAP (YAP+; *green*) and Hoechst (Hoechst+; *blue)* immunofluorescence in MIO-M1 after control or MAP4K4/6/7 siRNAs treatment.

(**H**) Quantification of relative YAP+ pixel intensity in Hoechst- MG area in (**G**).

(I) Quantification of relative YAP+ pixel intensity in Hoechst+ MG nuclei in (**G**).

For Western blots, levels were given as a.u. ± SEM in comparison with the control group (n = 3 independent pooled samples per group; Student’s *t*-test). **p*≤0.05, ***p*≤0.01, ****p*≤0.001.

**Figure S2. MAP4K4/6/7 inhibitor DMX-5804 suppresses YAP phosphorylation in MIO-M1.**

(**A, B**) Western blots (**A**) and quantification (**B**; normalized to vinculin) of p YAP and YAP in MIO-M1 after various concentrations of DMX-5804 treatment.

(**C, D**) Western blots (**C**) and quantification (**D**; normalized to vinculin) of p YAP and YAP in MIO-M1 at indicated time points after 5 μM DMX-5804 treatment.

(**E**) Immunofluorescence staining of YAP in MIO-M1 after 5 μM DMX-5804 treatment.

(**F**) Quantification of relative YAP+ pixel intensity in Hoechst- MG area in (**E**).

(**G**) Quantification of relative YAP+ pixel intensity in Hoechst+ MG nuclei in (**E**).

(**H-K**) Western blots (**H**) and quantification (**I** and **J**; normalized to vinculin) of p YAP and YAP after 6 h of various doses of DMX-5804 treatments in cells overexpressing empty vector or myc-MAP4K4. For Western blots, levels were given as a.u. ± SEM relative no-treatment (NT) (n = 3 independent pooled samples per group; Student’s *t*-test for **B,D** and **F**, one-way ANOVA for **J** and **K**). For pixel intensity measurements, levels were given as mean ± SEM (n = 3 per group; Student’s *t*-test). **p*≤0.05, ***p*≤0.01, ****p*≤0.001.

**Figure S3. MAP4K4/6/7 inhibitor DMX-5804 suppresses YAP phosphorylation in MG of murine retina.**

(**A, B**) Timeline diagram of the experimental procedures used in (**C-F**). Single intraperitoneal injection of DMX-5804 was carried out for concentration- (C) and time- (E) dependent test respectively. The *purple* triangles represent the time points of the sampling.

(**C, D**) Western blots (**C**) and quantification (**D**; normalized to vinculin) of p YAP and YAP after single dose of various concentrations of DMX-5804 injection.

(**E, F**) Western blots (**E**) and quantification (**F**; normalized to vinculin) of p YAP and YAP at indicated time points after 2 mg/kg DMX-5804 injection.

(**G**) YAP (YAP+, *red*) and SOX9 (SOX9+, *green*) immunofluorescence in mouse retinas after single DMX-5804 injection.

(**H**) Quantification of relative YAP+ pixel intensity in SOX9- MG nuclei in (**G**).

(**I**) Quantification of relative YAP+ pixel intensity in SOX9+ MG nuclei in (**G**).

For western blots, levels were given as a.u. ± SEM relative no-treatment (NT) (n = 3 independent pooled samples per group; Student’s *t* test). For pixel intensity measurements, levels are given as mean ± SEM (n = 3 per group; Student’s *t*-test). **p*≤0.05, ***p*≤0.01, ****p*≤0.001.

**Figure S4. MAP4K4/6/7 inhibitor DMX-5804 promotes the generation of MG-derived functional neurons.**

(**A**) The specificity of Glast-CreERT2+/tg;ROSA26R-tdTomato+/tg tracing system in mice.

Glast-CreERT2+/tg;ROSA26R-tdTomato+/tg mice were intraperitoneally injected with tamoxifen every 24 h for 5 consecutive days. Mice were sacrificed and whole flat-mounted retinas were prepared. The *purple* triangles represent the time points of the sampling.

(**B**) NeuN (*green*) immunofluorescence on whole flat-mounted mouse retinas after tamoxifen treatment.

(**C**) Timeline diagram of the experimental procedures used in (**D**). Glast-CreERT2+/tg;ROSA26R-tdTomato+/tg mice were intraperitoneally injected with Tamoxifen for 5 consecutive days. Then NMDA was intraocularly administered. DMX-5804 was injected every 6 h intraperitoneally from 6 hpi until 3 dpi after NMDA injection. The *purple* triangles represent the time points of the sampling.

(**D**) EdU labeling (*gray*) and NeuN (*green*) immunofluorescence in the NMDA-injured retinas after DMX-5804 treatment.

**Figure S5. The expression of proneural transcription factors in MG after NMDA/DMX-5804 induction.**

(**A**) Timeline diagram of the experimental procedures used in (**B-D**). Glast-CreERT2+/tg;ROSA26R-tdTomato+/tg mice were intraperitoneally injected with Tamoxifen for 5 consecutive days. Then NMDA was intraocularly administered. DMX-5804 was injected every 6 h intraperitoneally from 6 hpi until 5 dpi after NMDA injection. Then, 0,3,5,10,15 and 20 days after NMDA injection, mice were sacrificed and retinal samples were analyzed. The *purple* triangles represent the time points of the sampling.

(**B**) Heatmap of several key transcription factors in isolated MGs after NMDA/Solvent or NMDA/DMX-5804 treatment.

(**C** and **D**) Quantification of the mRNA levels of Ascl1 and Atoh7 in MG.

**Figure S6. The specificities of AAV tracing systems used in this study.**

(**A**) Diagram of the difference between GFAP and Short GFAP promotor in AAV vectors.

(**B**) Timeline diagram of the experimental procedures used in (**C, D**). Wildtype mice were intravitreally injected with pAAV-short GFAP-MCS-EGFP-3FLAG. Mice were sacrificed on 21^st^ day and whole flat-mounted retinas were prepared. The *purple* triangles represent the time points of the sampling.

(**C**) NeuN (*green*) immunofluorescence on whole flat-mounted retinas three weeks after infection with pAAV-short GFAP-MCS-EGFP-3FLAG.

(**D**) SOX9 (*red*) immunofluorescence on whole flat-mounted retinas three weeks after infection with pAAV-short GFAP-MCS-EGFP-3FLAG.

**Figure S7. Long-term tracing of MG fate after NMDA/DMX-5804 treatment by AAV tracing system.**

(**A**) Timeline diagram of the experimental procedures used in (**B-E**). Wildtype mice were intravitreally injected with pAAV-short GFAP-MCS-EGFP-3FLAG. NMDA was intraocularly administered 3 wk later. Then DMX-5804 was injected every 6 h intraperitoneally from 6 hpi until 5 dpi after NMDA injection. Thirty (30) days after NMDA injection, mice were sacrificed and the retinas were analyzed. The *purple* triangles represent the time points of the sampling.

(**B**) NeuN (NeuN+; *red*) immunofluorescence on whole flat-mounted NMDA-injured retinas retinal sections infected with pAAV-short GFAP-MCS-EGFP-3FLAG after solvent or DMX-5804 treatment.

(**C**) GABAergic cell marker GAD67 (GAD67+; *red*) or retinal ganglion cell marker (RBPMS+; *red*) immunofluorescence on whole flat-mounted retinas and retinal sections infected with pAAV-short GFAP-MCS-EGFP-3FLAG after NMDA/DMX-5804 treatment.

(**D**) Quantification of the number of NeuN+GFP+, GAD67+GFP+ cells or RBPMS+GFP+ cells per mm^2^ in (**B**) and (**C**).

(**E**) Quantification of the percentage of NeuN+GFP+, GAD67+GFP+ cells or RBPMS+GFP+ cells in GFP+ cells per mm^2^ in (**B**) and (**C**).

For quantification of NeuN+ GFP+ cells, levels were given as mean ± SEM (n = 3 retinas per group).

**Figure S8. Canonical Hippo pathway inhibitor XMU-MP-1 suppresses YAP phosphorylation in MG of murine retina.**

(**A,** **B**) Timeline diagram of the experimental procedures used in (**C-F**). Single intraperitoneal injection of XMU-MP-1 was carried out for concentration- (**C**) and time- (**E**) dependent test, respectively. The *purple* triangles represent the time points of the sampling.

(**C, D**) Western blots (**C**) and quantification (**D**; normalized to vinculin) of p YAP and YAP at 6 h after single dose of various concentrations of XMU-MP-1 injection.

(**E, F**) Western blots (**E**) and quantification (**F**; normalized to vinculin) of p YAP and YAP at indicated time points after single 2 mg/kg XMU-MP-1 injection.

(**G**) YAP (YAP+; *red*) and SOX9 (SOX9+; *green*) immunofluorescence on retinal sections at indicated time points after single 2 mg/kg XMU-MP-1 injection.

(**H**) Quantification of relative YAP+ pixel intensity in SOX9- MG nuclei in (**G**).

(**I**) Quantification of relative YAP+ pixel intensity in SOX9+ MG nuclei in (**G**).

For Western blots, levels were given as a.u. ± SEM relative no-treatment (NT) (n = 3 independent pooled samples per group; Student’s *t*-test). For pixel intensity measurements, levels were given as mean ± SEM (n = 3 per group; Student’s *t*-test ). **p*≤0.05, ***p*≤0.01, ****p*≤0.001.

**Figure S9.** **The comparison of abilities of XMU-MP-1 and DMX-5804 to promote MG proliferation and trans-differentiation.**

(**A**) Timeline diagram of the experimental procedures used in (**B-D**). Wild type mice were intravitreally injected with NMDA first. XMU-MP-1 or DMX-5804 was injected starting from 6 hpi until 5 dpi with injections every 6 h. EdU was injected intraperitoneally every 24 hours. Mice were sacrificed 5 days later. The red triangles represent the time points of the EdU injection and the *purple* triangles represent the time points of the sampling.

(**B**) EdU labeling (Edu+; *red*) and SOX9 (SOX9+; *green*) immunofluorescence on whole flat-mounted retinas 5 days after NMDA injection and consecutive XMU-MP-1 or DMX-5804 treatment.

(**C**) Quantification of the number of EdU+SOX9+ cells on whole flat-mounted retinas 5 days after NMDA injection and consecutive XMU-MP-1 or DMX-5804 treatment.

(**D**) Quantification of the percentage of EdU+ SOX9+ cells in SOX9+ cells on whole flat-mounted retinas 5 days after NMDA injection and consecutive XMU-MP-1 or DMX-5804 treatment.

(**E**) Timeline diagram of the experimental procedures used in (**F-H**). Two systems were used. One is Glast-CreERT2+/tg;ROSA26R-tdTomato+/tg mice were intraperitoneally injected with Tamoxifen, and the other is wild type mice were intravitreally injected with pAAV-short GFAP-MCS-EGFP-3FLAG. For Glast-CreERT2+/tg;ROSA26R-tdTomato+/tg mice which were injected with Tamoxifen, NMDA was administered into the vitreous 5 days later. For wild type mice, which were intravitreally injected with pAAV-short GFAP-MCS-EGFP-3FLAG, NMDA was administered into the vitreous 3 wk later. XMU-MP-1 or DMX-5804 was firstly injected starting from 6 hpi until 5 dpi with injections every 6 h. The *purple* triangles represent the time points of the sampling.

(**F**) TdTomato and NeuN (NeuN+; *green*) immunofluorescence or EGFP and NeuN (NeuN+; *red*) immunofluorescence on whole flat-mounted retinas after NMDA injection and consecutive XMU-MP-1 or DMX-5804 treatment.

(**G**) Quantification of the number of tdTomato+NeuN+ cells on whole flat-mounted retinas 30 days after NMDA injection and consecutive XMU-MP-1 or DMX-5804 treatment.

(**H**) Quantification of the percentage of tdTomato+NeuN+ cells in tdTomato+ cells on whole flat-mounted retinas 5 days after NMDA injection and consecutive XMU-MP-1 or DMX-5804 treatment.

(**I**) Quantification of the number of EGFP+NeuN+ cells on whole flat-mounted retinas 30 days after NMDA injection and consecutive XMU-MP-1 or DMX-5804 treatment.

(**J**) Quantification of the percentage of EGFP+NeuN+ cells in EGFP+ cells on whole flat-mounted retinas 5 days after NMDA injection and consecutive XMU-MP-1 or DMX-5804 treatment.

For quantification of EdU+ SOX9+, tdTomato+NeuN+ or EGFP+NeuN+ cells, levels were given as mean ± SEM (n = 3 retinas per group; Student’s *t*-test). **p*≤0.05, ***p*≤0.01, ****p*≤0.001.

**Figure S10. DMX-5804 but not XMU-MP-1 regulates JNK pathway activities in murine retina.**

(**A, B**) Western blots (**A**) and quantification (**B**; normalized to vinculin) of p JNK and JNK in murine retinas at 6 hours after a single dose of various concentrations of DMX-5804 injection.

(**C, D**) Western blots (**C**) and quantification (**D**; normalized to vinculin) of p JNK and JNK in murine retinas at 6 h after a single dose of various concentrations of XMU-MP-1 injection.

(**E, F**) Western blots (**E**) and quantification (**F**; normalized to vinculin) of p JNK and JNK at indicated time points after a single 2 mg/kg DMX-5804 injection.

(**G, H**) Western blots (**G**) and quantification (**H**; normalized to vinculin) of p JNK and JNK at indicated time points after a single 2 mg/kg XMU-MP-1 injection.

For Western blots, levels were given as a.u. ± SEM relative no-treatment (NT) (n = 3 independent pooled samples per group; Student’s *t-*test). **p*≤0.05, ***p*≤0.01, ****p*≤0.001.

**Figure S11. DMX-5804 suppresses JNK pathway activities in the NMDA-injured retina.**

(**A**) Timeline diagram of the experimental procedures used in (**B-G**). Wild type mice were intravitreally injected with PBS or NMDA. Solvent or DMX-5804 was injected starting from 6 hpi until 5 dpi with injections every 6 h. Mice were sacrificed at the indicated time points.

(**B-D**) Western blots analysis of p JNK and JNK at indicated time points after PBS (**B**), NMDA/solvent (**C**), or NMDA/DMX-5804 (**D**) injections. The *purpl*e triangles represent the time points of the sampling.

(**E-G**) Quantification (normalized to vinculin) of p JNK and JNK in (**B-D**).

For Western blots, levels were given as a.u. ± SEM in comparison with no-treatment (NT) group (n = 3 independent pooled samples per group; Student’s *t*-test). **p*≤0.05, ***p*≤0.01, ****p*≤0.001.

**Figure S12. DMX-5804 has higher potency than XMU-MP-1 to promote MG proliferation when drugs were treated after acute phase of retinal injury.**

(**A, B**) Western blots (**A**) and quantification (**B**; normalized to vinculin) of p MOB1 and MOB1 at indicated time points in the NMDA-injured retinas.

(**C**) Timeline diagram of the experimental procedures used in (**D-F**). Wild type mice were intravitreally injected with NMDA first. Next, XMU-MP-1 or DMX-5804 was injected starting from 6 hpi until 5 dpi with injections every 6 h. EdU was injected intraperitoneally every 24 h. Mice were sacrificed at day 5 post-retinal injury. The *red* triangles represent the time points of the EdU injection and the *purple* triangles represent the time points of the sampling.

(**D**) EdU labeling (EdU+; *red*) and SOX9 (SOX9+; *green*) immunofluorescence on whole flat-mounted retinas 5 days after NMDA injection and XMU-MP-1 or DMX-5804 consecutive treatment.

(**E**) Quantification of the number of EdU+SOX9+ cells on whole flat-mounted retinas 5 days after NMDA injection and XMU-MP-1 or DMX-5804 consecutive treatment.

(**F**) Quantification of the percentage of EdU+SOX9+ cells in SOX9+ cells on whole flat-mounted retinas 5 days after NMDA injection and XMU-MP-1 or DMX-5804 consecutive treatment.

For Western blots, levels were given as a.u. ± SEM in comparison with no-treatment (NT) group (n = 3 independent pooled samples per group; Student’s *t-*test). For quantification of EdU+ SOX9+ cells, levels were given as mean ± SEM (n = 3 retinas per group; Student’s *t-*test). **p*≤0.05, ***p*≤0.01, ****p*≤0.001.

**Figure S13. Working model of MAP4Ks inhibition promoted retinal regeneration.**

By using a selective small-molecule inhibitor, DMX-5804, which simultaneously inhibits MAP4K4, MAP4K6 and MAP4K7 to activate YAP in the retina, MGs are able to reprogram and transdifferentiate into retinal neurons expressing both amacrine and RGC markers after retinal damage in adult mice.
